# Interactors of sacsin’s DNAJ domain identify function in organellar transport and membrane composition relevant to ARSACS pathogenesis

**DOI:** 10.1101/2024.01.08.574743

**Authors:** Alexandre M. Paré, Zacharie Cheng-Boivin, Afrooz Dabbaghizadeh, Sandra Minotti, Heather D. Durham, Benoit J. Gentil

**Author notes:** Corresponding author B. J. Gentil, room 210, 475 Pine Avenue West, Room 210, Dep of Kinesiology and Physical Education, McGill University, Montreal, QC, H2W 1S4.

## Abstract

Autosomal Recessive Spastic Ataxia of the Charlevoix Saguenay (ARSACS) is caused by loss of function mutations in the *SACS* gene encoding sacsin, a 520kDa protein with multiple functional domains. The goal of this study was to identify client proteins interacting with the J domain, a cochaperone domain interacting with Hsp70 chaperones, to gain insights into sacsin’s function and its disruption in experimental models of ARSACS. Pull downs from mouse brain identified Rabs and Rab-associated proteins including Rab1b, ARF5 and endophilin B2, involved in organelle trafficking. In cell and mouse models of ARSACS, higher molecular weight species of Rab1b were identified on SDS-PAGE in addition to the normal 25kDa band and Rab1was retained in the soma along with membranous organelles (*i.e.*, ER, Golgi and ATG9 autophagic vesicles). These changes were reversed by expression of the DNAJ domain or the Ubl domain of sacsin and occurred independent of the formation of abnormal bundles of intermediate filaments, a key feature of ARSACS. Although Rab1b was associated with both Golgi and ER in both *Sacs*^+/+^ and *Sacs*^-/-^ cells, expression of the DNAJ domain or the Ubl domain of sacsin increased Rab1b association with ER and restored normal electrophoretic mobility. Finally, subcellular distribution of another membrane protein, neuroplastin, a key receptor for synapse formation and plasticity, also was impaired, pointing to a general problem in Rab-dependent membrane trafficking in the absence of sacsin.

## Introduction

Autosomal recessive spastic ataxia of the Charlevoix-Saguenay (ARSACS), caused by loss-of-function mutations in the *SACS* gene, has a high prevalence in the Charlevoix-Saguenay region, but occurs worldwide [1]. ARSACS is a progressive childhood-onset disease typically diagnosed around age six by the typical triad of cerebellar ataxia, peripheral neuropathy and pyramidal tract signs. It often is accompanied by ophthalmologic features including thickened, hypermyelinated retinal fibers and a thicker-than-average peripapillary retinal nerve fiber layer [2-5]. ARSACS patient-derived fibroblasts and mouse models present with altered mitochondrial network and morphology, intermediate filament bundling, dysregulated autophagic flux, and aberrant protein and organelle localization [6-11].

Sacsin is a gigantic protein (520kDa) that is ubiquitously expressed but enriched in the cytoplasm of neurons [12, 13]. The protein domains are: At the N-terminus, an ubiquitin-like domain (Ubl), which can bind to the proteasome 20Sα subunit [14]; three large sacsin repeat regions (SRR) termed SIRPT1, SIRPT2 and SIRPT3, which share homology with the chaperone, Hsp90; an XPC-binding domain, and at the C-terminus a J domain (SacsJ) homologous to Hsp40, immediately followed by a single higher eukaryote and prokaryote nucleotide-binding domain (HEPN) of unknown functions [15-18].

Evidence, including sequence homology to heat shock proteins and *in silico* analysis, points to various sacsin domains playing roles in protein chaperoning and quality control [18]. The XPC-binding domain of another protein, HR23A, binds to the ubiquitin ligase Ube3A, suggesting that sacsin might be ubiquitinylated by Ube3A[19]. The SSR region is specific to sacsin and exhibited *in vitro* chaperoning activity similar to HSP90 (*i.e.*; a “protective” chaperone activity) with additional “folding” chaperone action absent in Hsp90 [15, 16]. The SacsJ domain acted as a co-chaperone binding to Hsp70 chaperone family proteins to resolve aggregates made of GFP-ataxin-1[82Q] in SH-SY5Y cells [14]. Although the mechanism by which sacsin is involved in autophagy is not known, autophagy regulation was defective in SHSY5Y cells after knockout of the SACS gene and autophagic flux was increased in fibroblasts derived from ARSACS patients [11]. Finally, in addition to a role in the regulation of the cytoskeleton [20], recent omics approaches have identified a role of sacsin in intracellular transport of integrins and lysosomes [6, 7], calcium homeostasis [8], synaptic organization [9] and mitochondrial homeostasis [10, 11].

Recently, using STRING network analysis, sacsin was shown to interact with multiple heat-shock protein (HSP) chaperones, and to be a key element for trafficking to the plasma membrane of alpha integrins, which were sequestered into IF bundles [6]. We recently demonstrated direct interaction between IF and the sacsin J domain [21]; however, DNAJ domains typically have multiple clients [22]. Hence, we aimed to identify the SacsJ interactome to better understand the functions of sacsin and key elements of ARSACS pathogenic cascade.

## Materials and methods

### Plasmids and Recombinant protein production

Plasmids encoding wild type GFP-Rab1b GFP-Rab1bwt) were purchased from the MRC PPU reagent unit of University of Dundee (UK) cDNA Clone - Rab1b | MRC-PPU Reagents (dundee.ac.uk).

The DNAJ cDNA (corresponding to residues 4316-4420 of sacsin) was subcloned using mouse pEGFP-sacsin full length cDNA (OriGene Technologies, Rockville, MD, USA) and inserted in frame with GST into the pGEX6 vector. A myc epitope was tagged onto the C-terminus of pGEX-DNAJ or GST to identify the peptide after delivery into cells and tissues. The TAT-derived cell penetrating peptide sequence (YGRKKRRQRRR) [23], which has been shown to confer efficient neuronal delivery and blood-brain barrier penetration [24], was incorporated for delivery of recombinant polypeptide into cells. All cloning was subcontracted to NorClone (London, Ontario, Canada [NorClone - Gene Cloning Supplier](http://norclone.com/)).

For recombinant protein production, plasmids carrying the DNAJ-myc-TAT or GST-myc-TAT cDNA were transformed into Escherichia coli BL21 (DE3) pLysS. Bacterial cultures were grown overnight at 37°C and then diluted at 1:100 (v/v) into 1L Luria–Bertani medium containing 100μg/ml ampicillin. Bacteria were grown at 37°C with vigorous shaking at 225 rpm until they reach an OD600nm=0.6. Recombinant protein expression was then induced using 1mM isopropyl-1-thio-β-D galactopyranoside (IPTG) at 30°C for 4 hours.

Bacteria were lysed in PBS buffer (137 mM NaCl, 2.7 mM KCl, 10 mM Na_2_HPO_4_, 2 mM KH_2_PO_4_ pH 7.4), supplemented with 1 mM phenylmethylsulfonyl fluoride (PMSF), 80 units of DNase,100 µg/ml lysozyme and 0.1% Triton. After sonication, bacterial debris was pelleted using centrifugation. The supernatant was then passed through a 0.2 µm filter and chromatography affinity purification of GST-myc-TAT or GST-DNAJ-myc-TAT was carried out using glutathione resin according to the manufacturer instructions.

### Pull down and Proteomic analysis

Two mg of mouse brain were homogenized in Tris-buffered saline (TBS) (50 mM Tris-HCl, 150 mM NaCl, pH 7.5) 1% Triton X100 and centrifuged 3 times to remove debris and insoluble proteins. The sample was then incubated with 2 mg of recombinant GST or with GST-DNAJ with 200μl of Glutathione Sepharose resin for two hours at 4°C. At the end of the incubation, the suspension was centrifuged for 30 seconds at 5000xg at 4°C. Beads were washed 4–5 more times with 50mM ammonium bicarbonate prior to sending to the Proteomics Platform of the RI-MUHC, Montreal, Canada for further analysis (MUHC Proteomics Services - Research Institute of the McGill University Health Centre - RI-MUHC (rimuhc.ca)). Peptides were identified by MS/MS. Data were analysed using Scaffold4 software to identify specific partners of the DNAJ domain versus control GST; only proteins with more than two different peptides identified by Mass spectrometry (LS/MS) were considered as specific interacting partners. Quantitative analysis was performed on three samples per condition and statistical analysis was by T-test. STRING analysis of interactors (STRING: functional protein association networks (string-db.org)) was used to identify biological networks in which the DNAJ domain is involved.

### Cell culture

Immortalised fibroblasts were cultured in Dulbecco’s Modified Essential Medium with 10% fetal bovine serum (FBS). Control fibroblasts were from the CellBank Repository for Mutant Human Cell Strains (McGill University Health Complex, Montreal, QC, Canada). Fibroblasts were treated with GST-myc-TAT or GST-DNAJ-mycTAT to assess the time-dependence (30min to 24h) and dose-dependence (0-5μM) of the peptide’s effects on the vimentin network, evaluated by immunolabeling with anti-vimentin antibodies (clone V9, MA5-11883 Thermofisher). Peptides were identified by indirect immunofluorescence using an anti-myc antibody (C3956, Sigma-Aldrich). SW13-human adrenocarcinoma vimentin-deficient cells, lacking intermediate filaments, were grown in DMEM + 10% FBS [25]. SW13-cells lacking sacsin expression (SW13-*^SACS^*^-/-^) were produced using the sacsin double nickase plasmid following the manufacturer’s instructions (SantaCruz Biotechnology, cat. no. sc-404592-NIC).

Primary cultures of dissociated spinal cord-DRG were prepared from E13 *Sacs^+/+^* and *Sacs^-/-^*mice (C57Bl6 background). Generation and characterization of the *Sacs^-/-^*mice were as previously described (Lariviere et al., 2015); wild type (*Sacs^+/+^)* mice on the same background were used as control (*Sacs^+/+^*). Cultures were plated on glass coverslips (Fisher, ON, Toronto, Canada) coated with poly-D-lysine (P7280, Sigma-Aldrich) and Matrigel® (CACB354234, VWR, Town of Mount Royal, QC, Canada) and maintained in Eagle’s Minimum Essential Medium enriched with 5 g/l glucose and supplemented with 3% horse serum, and other growth factors as previously described [26]. Cultures were used in experiments 6 weeks following plating to allow neuronal maturation and apparition of neurofilament bundles in more than 80% of *Sacs^-/-^*motor neurons. Neurofilament bundles were defined as previously by a continuous bundle crossing the cell body from dendrite to dendrite [25] in neurons immunolabeled with anti-NFL (clone NR4, N5139, Sigma-Aldrich).

### Processing of mouse brain and spinal cord

Paraformaldehyde-fixed brain and spinal cord of 7 month-old *Sacs^+/+^*and *Sacs^-/-^* mice (C57Bl6 background) were processed at the Histopathology - Research Institute of the McGill University Health Centre - RI-MUHC (rimuhc.ca) for paraffin embedding and sectioning (10μm thin). The sections were then deparaffinised to carry out indirect immunofluorescence labelling.

### Antibodies for immunoblotting

Primary antibodies used for immunoblotting were: mouse monoclonal anti-NFL antibody clone NR4 (N5139, Sigma-Aldrich, 1:400), mouse monoclonal anti-flag antibody clone M2 (F3165, Sigma-Aldrich, 1:400), rabbit polyclonal anti-Rab1b (SPA810, Enzo Life Sciences, Framingdale, NY, USA), rabbit polyclonal anti-endophilinB2 (15897-1-AP, Proteintech, USA, 1:400), rabbit polyclonal anti-ARF5 (clone 9E10, Santa Cruz Biotechnology, Santa Cruz, CA, USA, 1:400), rabbit polyclonal anti-ATG9A (26276-1-AP, Proteintech, 1:300), rabbit polyclonal anti-GM130 (11308-1-AP, Proteintech, 1:300), rabbit polyclonal anti-ERGIC53 (13364-1ap, Protein tech, 1:300), mouse monoclonal anti-myc (A14, Santa Cruz Biotechnology, 1:400), and mouse monoclonal anti-GAPDH, (MM-0163-P, MediMabs, 1:1,000), mouse monoclonal anti-vimentin (V6639, Sigma-Aldrich, 1:300), DAPI (1:1000), rabbit monoclonal anti-sacsin (ab181190-100UL, Abcam, 1:1000), rabbit monoclonal anti-human/mouse neuroplastin (AF5360, RNDsystems, 1:1000), rabbit polyclonal anti-Rab7a (55469-1-AP, Proteintech, 1:1000), mouse monoclonal anti-Rab5a (clone 1B6A5, Proteintech, 1:1000), rabbit anti-Rab1b antibody (Cat # 17824-1-AP Proteintech 1:1000).

Secondary antibodies were: Peroxidase-conjugated affinipure donkey anti-mouse and anti-rabbit IgG (715-035-150 and 711-035-152, 1:5,000), Cy2-conjugated (715-225-151, 1:300) and Cy3-conjugated (715-165-151, 1:300) donkey anti-mouse IgG and Cy3-conjugated (705-165-147, 1:300) donkey anti-goat IgG were from Jackson ImmunoResearch Laboratories (West Grove, PA, USA), Alexa Fluor 488-conjugated rabbit polyclonal (A-11034, Mol Probes, 1:300) and horseradish peroxidase (HRP)-conjugated goat anti-rabbit IgG (Proteintech, Cat # SA00001-2)

### Western analysis

Protein samples were separated by 10-12% SDS-PAGE and electrotransferred to nitrocellulose membranes (Merck Millipore, Cat # ISEQ00010). Immunoblotting was carried out as follows. After blocking with 5% milk, the membranes were incubated with the primary antibody, rabbit anti-Rab1b or rabbit anti-sacsin, at 4°C for 12h. The membranes were then incubated with HRP-conjugated goat anti-rabbit IgG secondary antibody (1:2500) for 2h. Finally, the signal was detected using enhanced chemiluminescence (ECL, Thermofisher) and images were acquired with Intas Chemiluminescence imager (Science imaging, Sweden); blots were reprobed with mouse anti-GAPDH and HRP-conjugated goat anti-mouse as loading control.

### Statistical analysis of IF and organelle organization

Quantitative experiments are presented as mean ± SD. The proportion of motor neurons or fibroblasts carrying a dismantled, regular or perinuclear IF network was quantified in at least three coverslips per condition, with a minimum count of 30 cells per coverslip.

The proportion of motor neurons displaying a widely distributed or dendritic-, or somal-restricted organelle distribution was quantified using at least three coverslips per condition, with a minimum count of 30 motor neurons per coverslip. Statistical analysis was by one-way ANOVA followed by a Tuckey HSD post hoc analysis. P values < 0.05 were considered statistically significant.

### Colocalization analysis

Colocalization measurement is defined by the Pearson Correlation coefficient (PCC), which is a reliable and simple measure to evaluate proximity of fluorescent probes [27]. Rab1b colocalization with Golgi or ER was assessed by confocal microscopy following co-transfection of GFP-Rab1bwt and sacsin domains in SW13- and SW13^-SACS-/-^ cells, and indirect immunofluorescence double labelling with antibodies against the myc epitope and GM130 or KDEL, markers of Golgi and ER, respectively. PCC was measured using the ZenBlue 2.6 colocalization software tool. Statistical analysis was performed by one-way ANOVA followed by a Tuckey HSD posthoc analysis to determine significance for colocalization coefficients. P values < 0.05 were considered statistically significant. Three individual coverslips for a total of at least 30 cells per condition were analyzed.

## Results

### Identification of new partners of Sacsin J domains

DNAJ client proteins were identified in mouse brain extracts by a pulldown assay using recombinant sacsin DNAJ domain or GST as control. Proteins were sequenced by mass spectroscopy and quantified as proteins specific to or statistically enriched in the sacsin DNAJ data set relative to GST, shown as green squares or triangles in Volcano plots (Fig. 1A and 1B). A list of statistically significant interacting partners is displayed in the table of Fig. 1D, ordered by most enriched (top) to least enriched (bottom) proteins. Consistent with previous data, IF proteins were found in the interactome, which validated our approach. Novel specific partners including Rab1b are highlighted in yellow (Fig. 1C). Next, the STRING interaction database was used to generate a map of protein-protein interactions according to their function in cellular processes (Fig. 1E). Analysis of Enrichment of Gene Ontology identified regulation of subcellular localization of organelles and protein (53%), Golgi and vesicle transport (19%), and antigen processing and presentation of peptides (12%) as major cellular pathways involving DNAJ domain interactors (Fig. 1F).

**Figure 1.**
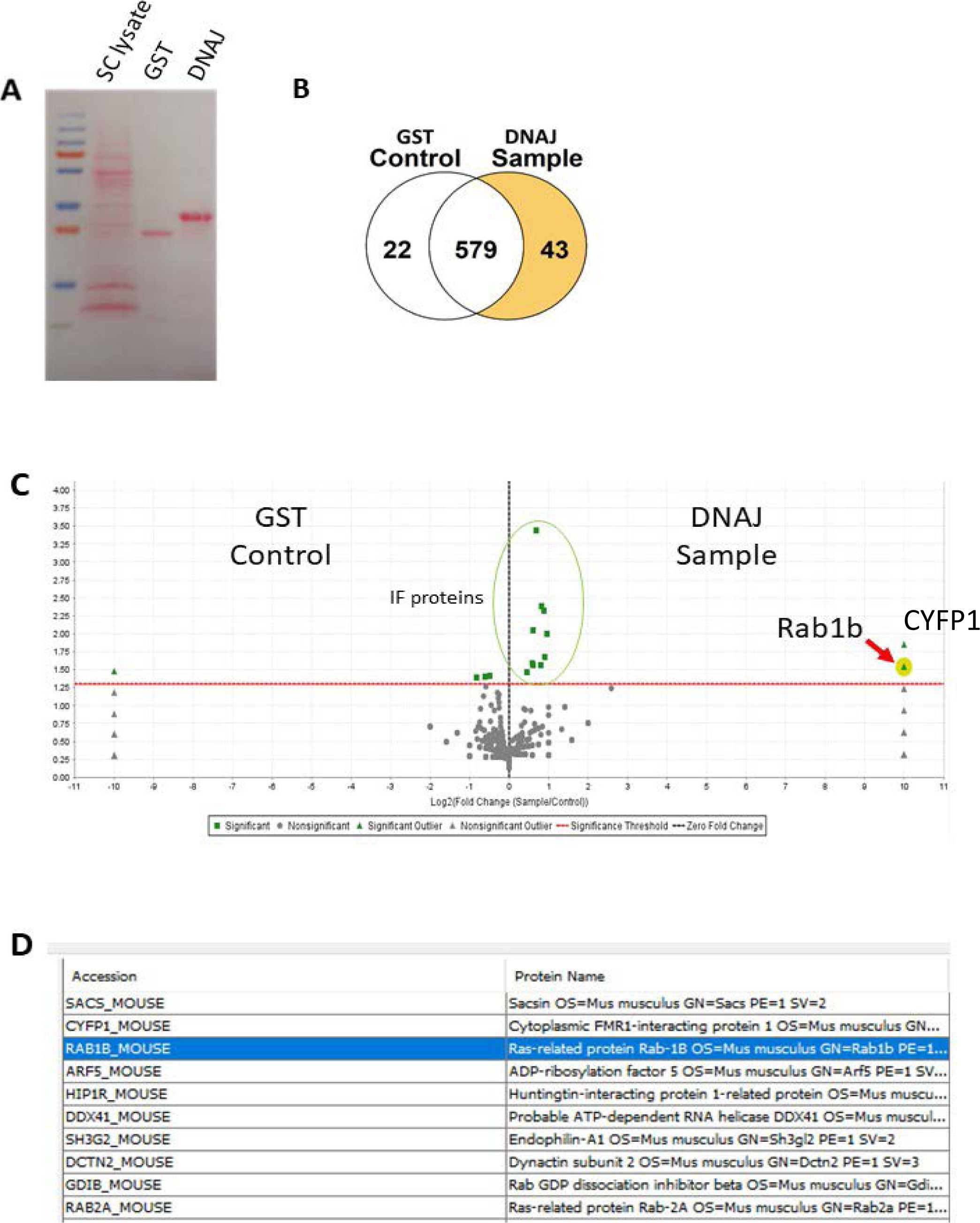
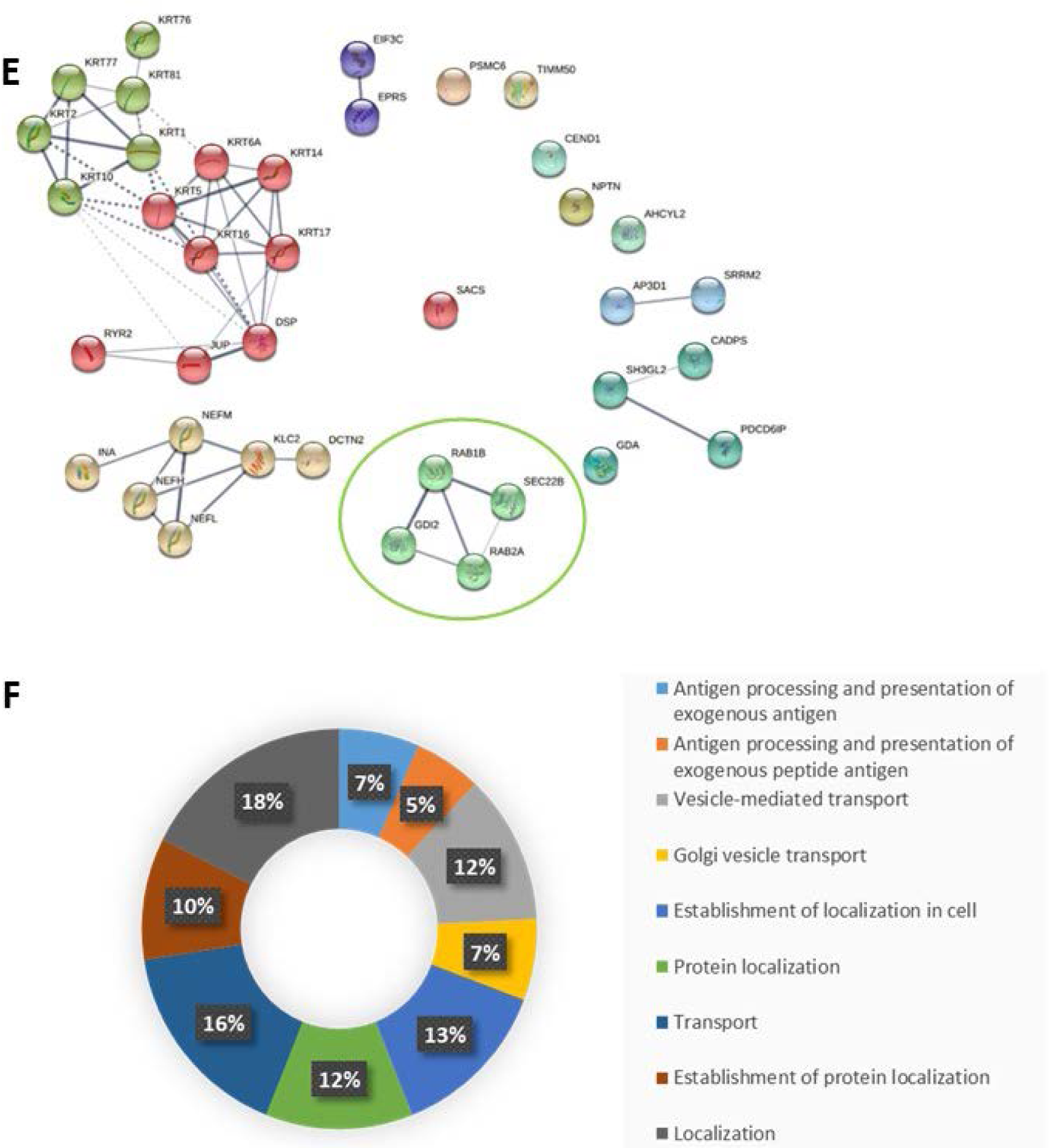
The interactome of the J domain of sacsin in mouse brain extracts showing associations with organellar localization and transport. **A)** Red ponceau staining of a nitrocellulose membrane following electrotransfer of SDS-PAGE. First lane: molecular weight markers; second lane: total spinal cord lysate; third lane: GST pulldown sample, and fourth lane: DNAJ pulldown. **B)** Number of proteins identified by mass spectrometry in the DNAJ and control GST pulldowns. using the Scaffold software. **C)** Volcano plot showing old Change of major proteins identified by mass spectrometry. Rab1b is significantly enriched in the DNAJ sample and is highlighted in yellow. **D)** Table listing identified major novel interactors, other than IF proteins, from the most enriched proteins, starting from the most enriched (top. Rab1b is second most enriched DNAJ-interacting protein. **E)** Interaction map of the DNAJ-specific partners using STRING software. DNAJ-interacting proteins were grouped by known functional networks. A cluster containing Rab1b with a PPI enrichment value of 0.00515 was identified (circle in green). **F)** Donut Pie-chart indicating major biological functions associated with DNAJ-interacting proteins as defined by GO-analysis. Organellar localization and transport were major functions identified.

Amongst all the DNAJ interacting partners, Rab1b was highly specific and was the second most enriched protein, with a function in organelle dynamics and autophagy, processes known to be disrupted in ARSACS. Next, the relevance of Rab1b in our interactome was validated by investigating expression levels and subcellular localisation in *Sacs^+/+^* and *Sacs^-/-^* mouse spinal cord. Western blot analysis of mouse spinal cord extracts identified an overall upregulation of Rab1b in the absence of sacsin and a different migration pattern of Rab1 on SDS-PAGE, with an additional band of lower electrophoretic mobility suggestive of post-translational modification (Fig. 2A). In addition, Rab1b was concentrated in the soma of *Sacs^-/-^* motor neuron in spinal cord (Fig. 2B); a similar pattern was observed *in vivo* in Purkinje neurons (supplemental S1).

**Figure 2.**
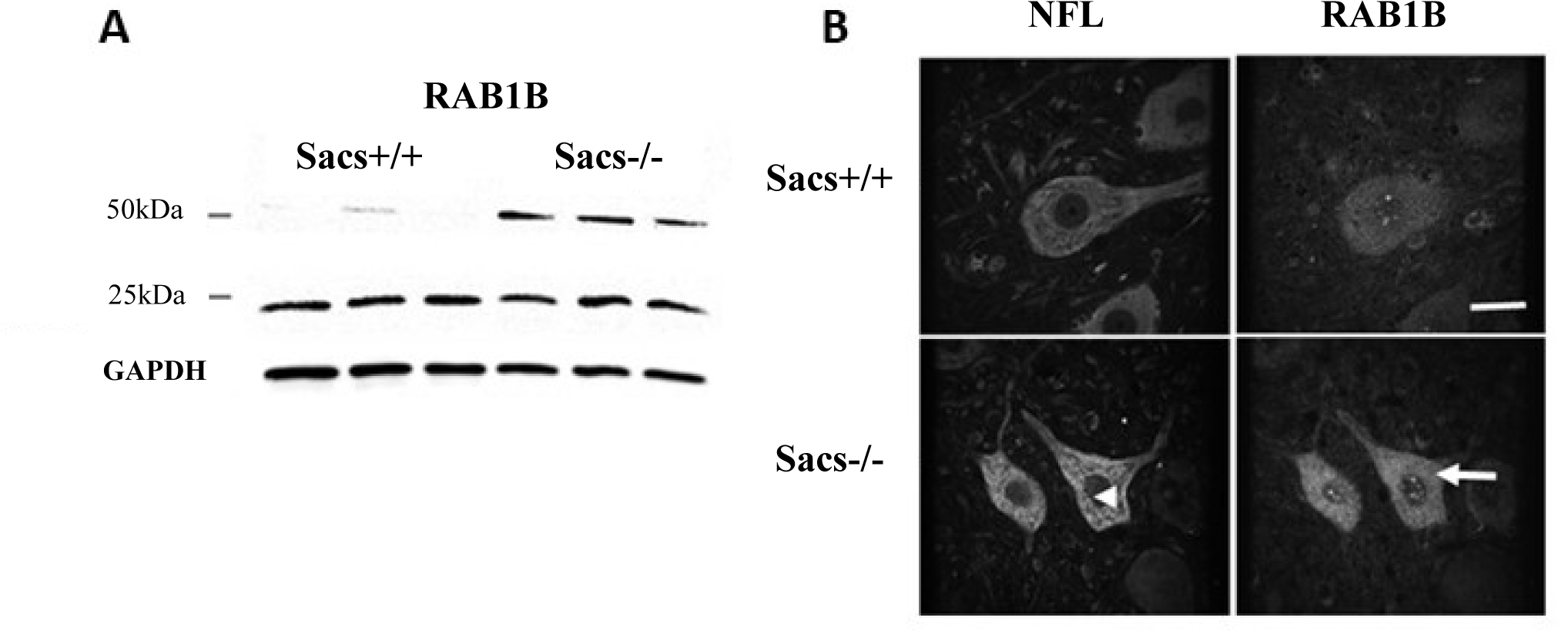
Altered processing of Rab1b and restricted distribution in *Sacs* -/- motor neurons. **A)** Western blot analysis of Rab1b in extracts of spinal cord from 6 month-old *Sacs^+/+^*and *Sacs^-/-^* mice. Rab1b was upregulated in *Sacs^-/-^*spinal cord and showed additional bands of higher molecular weight. **B)** Representative Z-projection of confocal images of indirect double immunofluorescence labelling of Rab1b (rabbit anti-rab1b) and NFL (mouse anti-NFL) in motor neurons of 6-months old *Sacs^+/+^* and *Sacs^-/-^* mouse spinal cord, showing NFL bundles and increased expression of Rab1b in *Sacs^-/-^* tissue.

### Sacsin plays a role in subcellular distribution of ER and Golgi and in autophagy through Rab1b

As shown in Fig. 3A, Rab1b is a key small GTPase regulating membrane dynamics between ER-Golgi intermediate compartment (ERGIC) or autophagic membranes labelled by ATG9.We next confirmed the abnormal distribution of Rab1b in cultured motor neurons and investigated the consequence of the lack of sacsin on Rab1b-dependent organelles. Subcellular distribution of Rab1b in cultured 6-week-old *Sacs^+/+^* and *Sacs^-/-^* spinal motor neurons was similar to the localisation of Rab1b in motor neurons *in vivo*. In cultured *Sacs^+/+^* murine motor neurons, Rab1b immunolabeling was widely distributed including in the distal dendrites, whereas in *Sacs^-/-^* motor neurons, Rab1b was abnormally restricted to the soma and absent from distal dendrites (Fig. 3B). While Rab1b-interacting organelles, ER, ERGIC, and Golgi are usually widely distributed and reach the extremities of distal dendrites in healthy motor neurons, they were restricted to the soma and absent in the distal dendrites in *Sacs^-/-^* motor neurons. This distribution pattern was consistent with the abnormal localisation of Rab1b (Fig. 3C-E). In addition, ATG9-positive autophagic membranes are usually found widely distributed in the soma and present in proximal dendrites of *Sacs^+/+^*motor neurons (Fig. 3F); however, they were abnormally compacted in the soma in *Sacs^-/-^* motor neurons and ATG9 immunofluorescence intensity was higher (Fig. 3F), which is consistent with involvement of sacsin in autophagy. Qualitatively, ATG9 membranes were juxtaposed to NF bundles, which suggests that NF play a role in anchoring autophagosomes (Fig. 3G).

**Figure 3.**
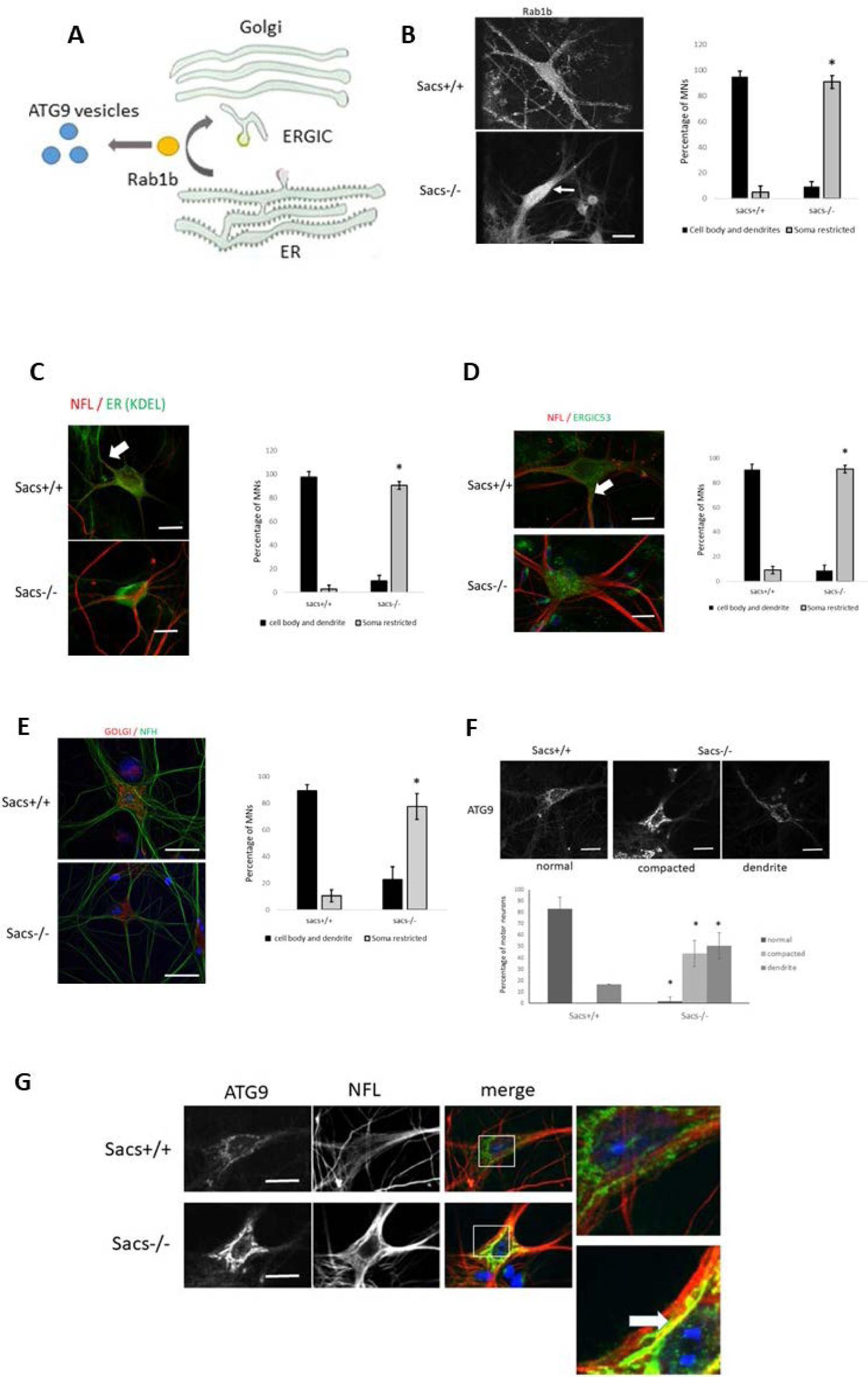
Distribution of Rab1b, ER, ERGIC, Golgi, and ATG9 is restricted in *Sacs^-/-^*motor neurons. **A)** Schematic representation of the role of Rab1b in the secretory pathway and autophagy. Rab1b plays a role in membrane fission, transport, and fusion at the ER, ERGIC, Golgi, and early steps of autophagy. **B)** Representative Z-projection of confocal images of indirect immunofluorescence labelling of Rab1b (rabbit anti-rab1b) and NFL (mouse anti-NFL) in 6-week-old spinal cord *Sacs^+/+^* and *Sacs^-/-^*motor neurons in culture showing accumulation of Rab1b in the soma of *Sacs^-/-^*motor neurons (bottom). Quantitation of the percentage of motor neurons showing distribution of Rab1b in cell body and dendrites versus restricted to the soma; *p< 0.05 vs *Sacs^+/+^* by one-way ANOVA, HSD Tukey post hoc analysis (N=3). Scale bar = 20μm. **C-G)** Distribution of membranous organelles in motor neurons of dissociated spinal cord cultures prepared from E13 *Sacs^+/+^* and *Sacs^-/-^* mouse embryos, maintained in culture for six weeks. Shown are representative Z-stacks of confocal images following indirect double immunofluorescence labelling with antibodies to organelle biomarkers and quantitation of biomarker according to distribution in the cell body and dendrites or restricted to the soma. C) The ER marker KDEL and NFL; D) ERGIC53 and NFL; E) the Golgi marker GMF130 and NFH; F) the autophagy marker ATG9 and NFL. *p< 0.05 vs *Sacs^+/+^* using a one-way ANOVA, HSD Tukey post hoc analysis. **G)** ATG9 was juxtaposed to NFL bundles (thick white arrow).

### Loss of sacsin alters Rab1b and organelle distribution independent of IF bundling

IF including NF are well known to have secondary effects on morphology and subcellular distribution of membranous organelles. Thus, we questioned whether the restricted distribution of Golgi, ER and autophagic vacuoles could be secondary to the formation of NF bundles in cells lacking sacsin. Rab1 expression and organelle distribution was evaluated in adrenocarcinoma SW13-cells lacking endogenous intermediate filaments. Like in cultured *Sacs^-/-^*murine motor neurons, Rab1b was upregulated and an extra band of higher molecular weight was observed on immunoblots of SW13-^SACS-/-^ cells (Fig. 4A). Abnormal Golgi morphology (*i.e.,* compaction and restriction of subcellular localization) also occurred in SW13-*^SACS^*^-/-^ (Fig. 4B), demonstrating that Rab1b and Golgi abnormalities in absence of sacsin expression were independent of IF bundles. To rule out nonspecific interactions of Rab1b antibodies, Rab1b was overexpressed in SW13-cells, with or without sacsin knockout, to determine if the higher molecular species observed on Western blots were specific. In SW13- ^SACS-/-^ cells, ectopic expression of flag-tagged Rab1 presented the same higher molecular species. These higher molecular bands were minor forms compared to the major 25kDa band (Fig. 4C). All together, the data show that Rab1b upregulation and abnormal post-translational modification and abnormal Golgi compaction are specific to the lack of sacsin, not a secondary effect of IF bundling.

**Figure 4.** The Golgi apparatus is compacted in SW13^- *SACS*-/-^ cells lacking endogenous intermediate filaments. **A)** Western blot analysis showing the upregulation of endogenous monomeric Rab1b (arrow) with higher weight molecular bands in SW13^-^ *^SACS^*^-/-^ cells, absence of sacsin expression in SW13^-SACS-/-^ cells and GAPDH as a loading control. **B)** Representative Z-stack confocal images following indirect immunofluorescence labelling for the Golgi marker GM130 and DAPI in SW13- and SW13^-^ *^SACS^*^-/-^ cultured cells. Quantitative assessment of the area of Golgi distribution by the total number of pixels of anti-GM130 labelling illustrates a compacted Golgi in the majority of SW13^-^ *^SACS^*^-/-^ cells. *p< 0.05 vs *Sacs^+/+^* using a one-way ANOVA, HSD Tukey post hoc analysis. Scale bar = 20μm.**C)** Western blot analysis of SW13- and SW13- *^SACS^*^-/-^ cells expressing Rab1b-flag showing a similar migration pattern as endogenous Rab1b and the high molecular weight forms in SW13- *^SACS^*^-/-^ samples.

### Sacsin domains alter Rab1b subcellular distribution and association with ER in SW13-*^SACS^*^-/-^ cells

Rab1b is required for vesicle transport between the ER and the Golgi [28]. Given the unbalance of Rab1b expression and compaction of the Golgi in cells lacking sacsin, Rab1b association with Golgi or ER was characterized by measuring the PCC of ectopically expressed Rab1b-EGFP (to avoid nonspecific Rab1 antibody labeling) with GM130 or KDEL, markers of Golgi and ER, respectively. EGFP-Rab1b was generally enriched at the Golgi, like endogenous Rab1b visualized by immunocytochemistry, and was strongly co-localized with GM130 in SW13- and SW13^-SACS-/-^ cells (PCC values of 0.87 and 0.84, Fig. 5A). EGFP-Rab1b also localized at the ER membrane, but to a lesser extent (PCC values of 0.25 and 0.28) (Fig. 6A). Expression of the DNAJ domain or other sacsin domains did not significantly alter Rab1b-EGFP colocalization with the Golgi apparatus in either SW13- and SW13^-SACS-/-^ cells (Fig. 5B-C). However, expression of the DNAJ, as well as DNAJH33Q (a variant that lacks ability to bind HSP70), and Ubl sacsin domains significantly increased Rab1b association with the ER, whereas SIRPT1 and HEPN domains had no effect (Fig.6B-C). The data demonstrate that ER enrichment is not dependent on HSP70 binding, SIRPT1 or HEPN functions.

**Figure 5.**
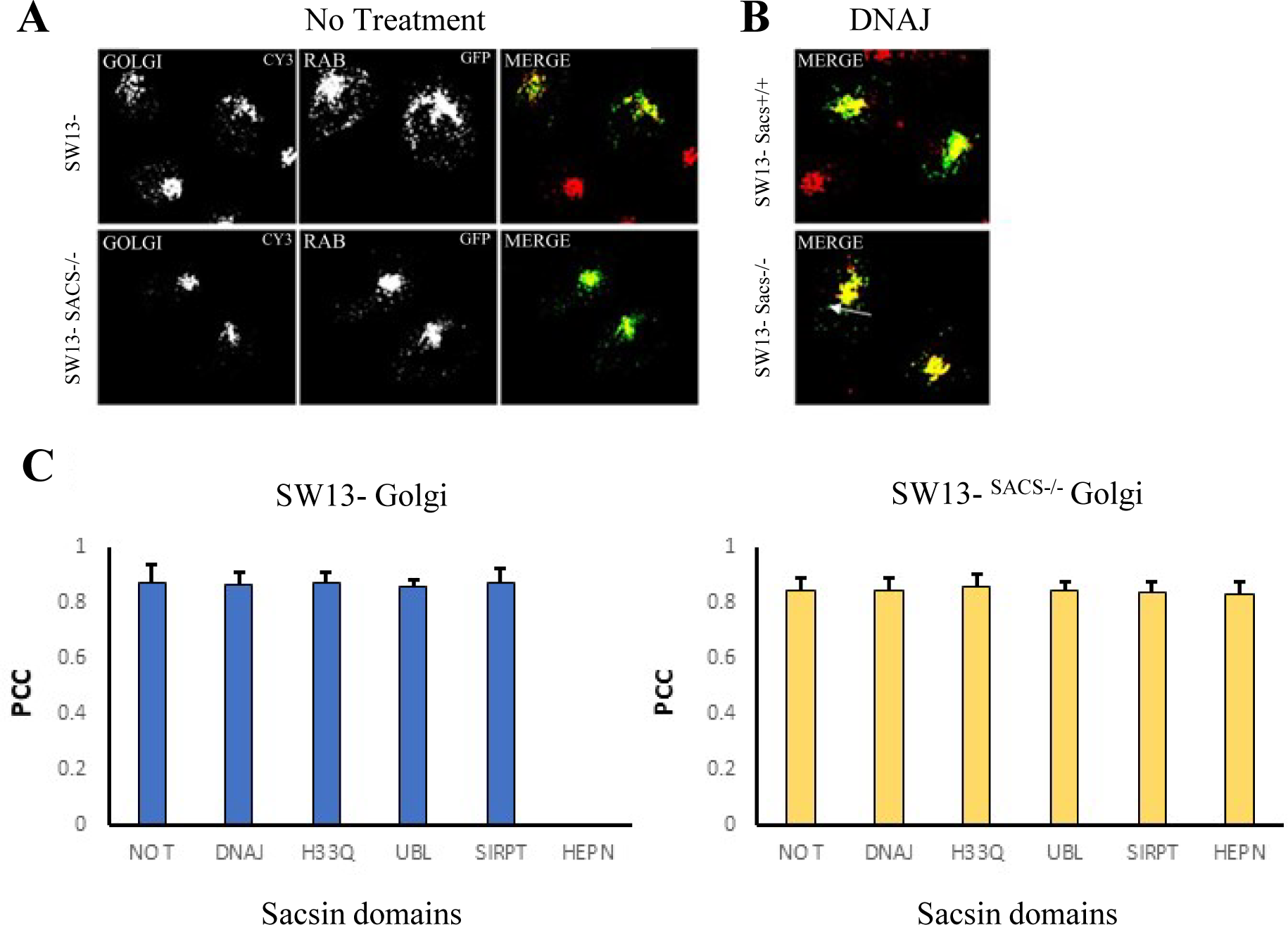
Localization of Rab1b. **A)** Representative Z-stacks of confocal images of indirect double immunofluorescence labelling for Rab1 and GM130 to assess colocalization of Rab1b and Golgi apparatus in SW13-*^SACS^*^+/+^ and SW13^-^ *^SACS^*^-/-^ cells following co-transfection of GFP-Rab1bwt and plasmids encoding individual sacsin domains. Yellow illustrates overlap of labelled structures in merge. **B)** Effect of the DNAJ domain on Rab1b colocalization with Golgi apparatus in SW13- and SW13^-^ *^SACS^*^-/-^ cells. **C)** Quantitation of colocalization between Rab1b and the Golgi apparatus following expression of various domains. A greater Pearson Correlation Coefficient (**PCC**) indicates greater colocalization of GFP-Rab1bwt with GM130. *p< 0.05 vs No Treatment counterparts using a one-way ANOVA, HSD Tukey post hoc analysis. Scale bar = 20μm. (N=25 per condition).

**Figure 6.**
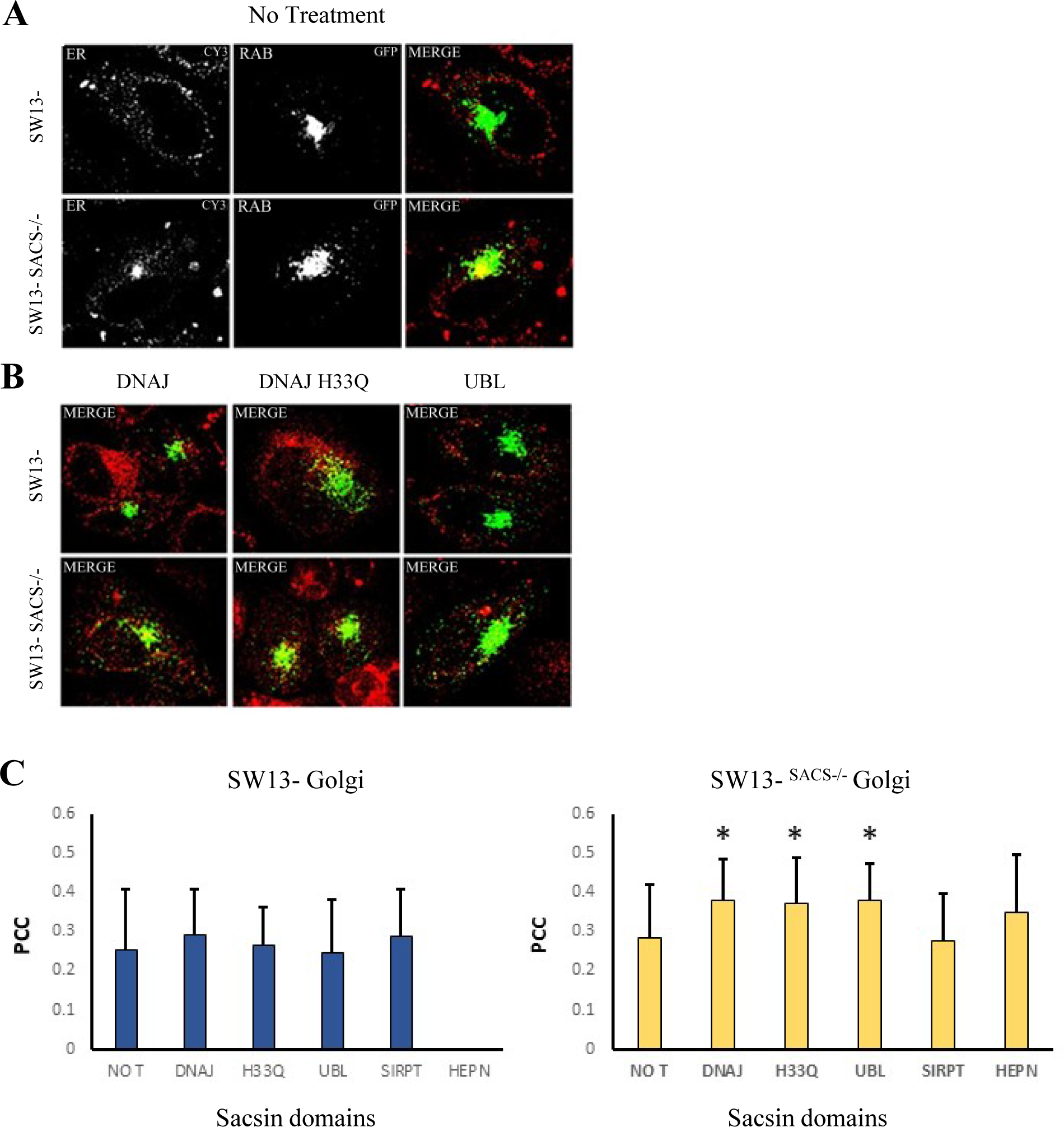
Rab1b is weakly co-localized with the ER and the sacsin domains DNAJ, DNAJ^H33Q^ (an HSP70-binding incompetent variant), and Ubl significantly increased Rab1b association with the ER in SW13-*^SACS-/-^* cells. **A, B)** Representative Z-stack of confocal images of indirect immunofluorescence labelling for the ER marker KDEL (red) in SW13*-* and SW13^-^ *^SACS^*^-/-^ cells after transfection with plasmid encoding GFP-Rab1b^wt^ A) alone or B) with plasmid encoding the sacsin domain DNAJ, DNAJH33Q (an HSP70-binding incompetent mutant) or Ubl and yellow illustrates overlap of labelled structures in merge. **C)** Quantitation of colocalization between Rab1b and ER following transfection of plasmid encoding domains. A greater Pearson Correlation Coefficient (PCC) indicates greater colocalization of GFP-Rab1bwt with KDEL. *p< 0.05 vs No Treatment counterparts using a one-way ANOVA, HSD Tukey post hoc analysis. Scale bar = 20μm. (N=30 per condition).

We next determined if expression of individual sascin domains could alter the expression profile of Rab1 and modify the higher molecular weight species of Rab1b in sacsin knockout cells (Fig. 7A). The higher molecular weight species of endogenous Rab1b was reduced by expression of the DNAJ domain and eliminated by expression of the Ubl domain. A similar pattern was observed with Rab5a and Rab7a, suggesting a general effect on Rabs processing (Fig. 7B-C). Because of the poor detergent solubility, we have not yet succeeded in identifying post-translational modifications responsible for the increased apparent molecular weight by mass spectrometry (Fig. 7).

**Figure 7.**
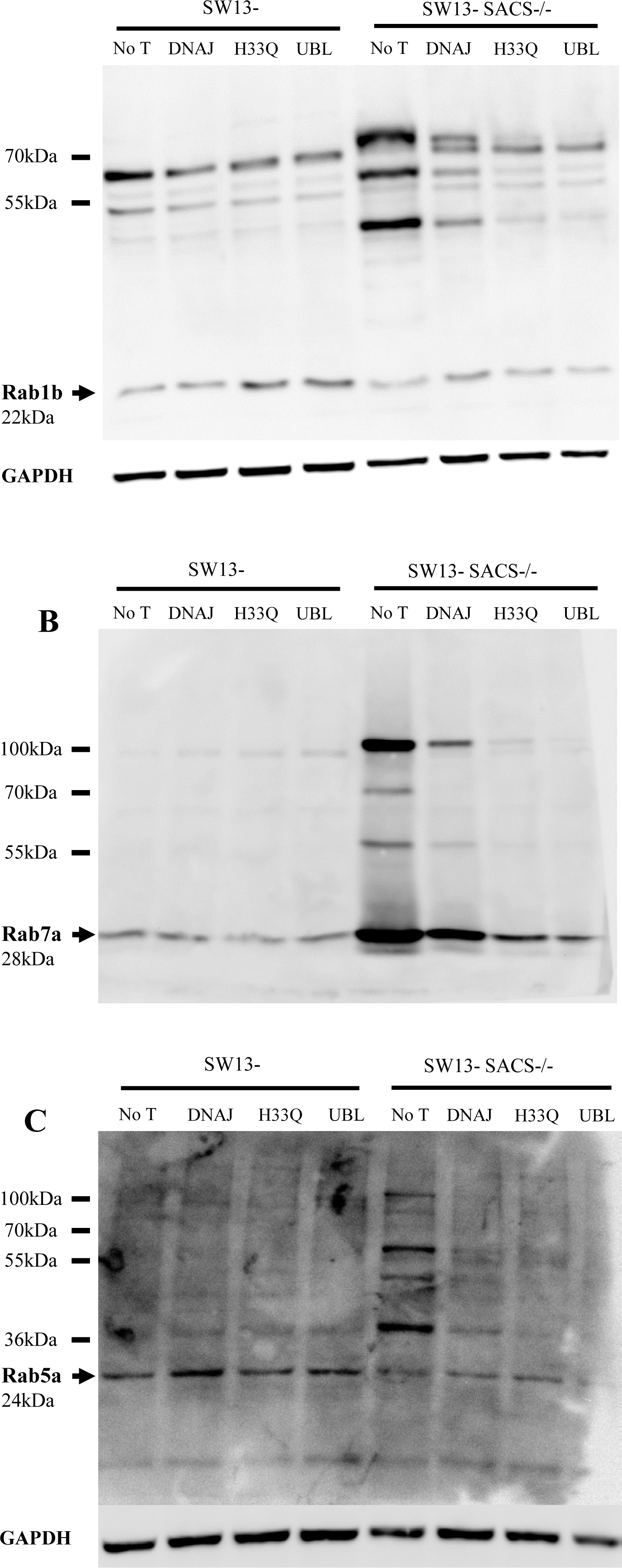
Upregulation of Rabs and different electrophoretic mobility patterns in SW13-*^SACS-/-^* cells by expression of sacsin DNAJ Ubl domain. **A)** Rab1b was upregulated and presented with additional higher molecular weight bands on Western blots of SW13- *^SACS-/-^* cells. Transfection of plasmids encoding these sacsin domains reversed the Rab1b expression pattern to wild-type. SDS-PAGE/Western blot analysis of Rab1b protein expression in SW13- and SW13^-^ *^SACS^*^-/-^ cells following expression of sacsin DNAJ, DNAJH33Q (Hsp70 binding incompetent) or Ubl domain. GAPDH served as a control for protein loading. A similar behavior was observed for **B**) Rab7a **C**) Rab5a.

In addition to Rab1b, Endophilin B2 and ARF5, both regulators of intracellular membrane dynamics, were identified in our interactome and reinforced a direct role of sacsin in organellar dynamics. (Subcellular distribution of Endophilin B2 was affected in cultured *Sacs^-/-^* 6-months-old of *Sacs^+/+^* mouse spinal motor neurons Fig. S3C). ARF5 accumulated in the soma of *Sacs^-/-^* motor neurons in culture as shown by the increased fluorescence intensity of ARF5 in the soma of *Sacs^-/-^* motor neurons compared to wild-type motor neurons (Fig. S3D). Altogether, these results support sacsin playing a crucial role in regulating membrane trafficking through interactions with Rabs and Rab-associated proteins.

### Sacsin plays a role in neuroplastin trafficking to the plasma membrane

We next examined whether Rab abnormalities and disruption of vesicle trafficking would alter export and expression of membrane proteins in *Sacs*^-/-^ cells. We focused on Neuroplastin (NPTN), which also was identified in the pulldown experiment, but by only two peptides. NPTN is a glycosylated cell adhesion molecule of the immunoglobulins superfamily that plays a role in synaptic plasticity and inhibits the internalization and desensitization of the GABA receptor. On Western blots, NPTN was labeled as a major band at 55 kDa and a minor band at 65 kDa in SW13-cells, corresponding respectively to the N-glycosylated NPTN65 and NPTN55 isoforms [29]. Glycosylated NPTN65 was absent in SW13^-SACS-/-^ (Fig. 8A) while lower forms at 40kDa and 30kDa were present, which could correspond to molecular weight of the non-glycosylated forms [30] and suggests altered processing, in particular absence of N-glycosylation. NPTN was widely distributed through the SW13-cells whereas in SW13^-SACS-/-^ cells, NPTN accumulated in the same region of the cell as the compacted Golgi apparatus (Fig. 8B), suggesting that lack of sacsin impacted the subcellular distribution of NPTN possibly through impairment of its transfer to the plasma membrane and defective maturation (*i.e.*; N-glycosylation). In *Sacs^+/+^* motor neurons, NPTN was concentrated at the plasma membrane with a distribution consistent with synaptic localization, whereas in *Sacs^-/-^*cultured motor neurons, NPTN was localized in the soma rather than the plasma membrane (Fig. 8C). A similar pattern was observed in Purkinje neurons *in vivo* (Fig. 8D)

**Figure 8.**
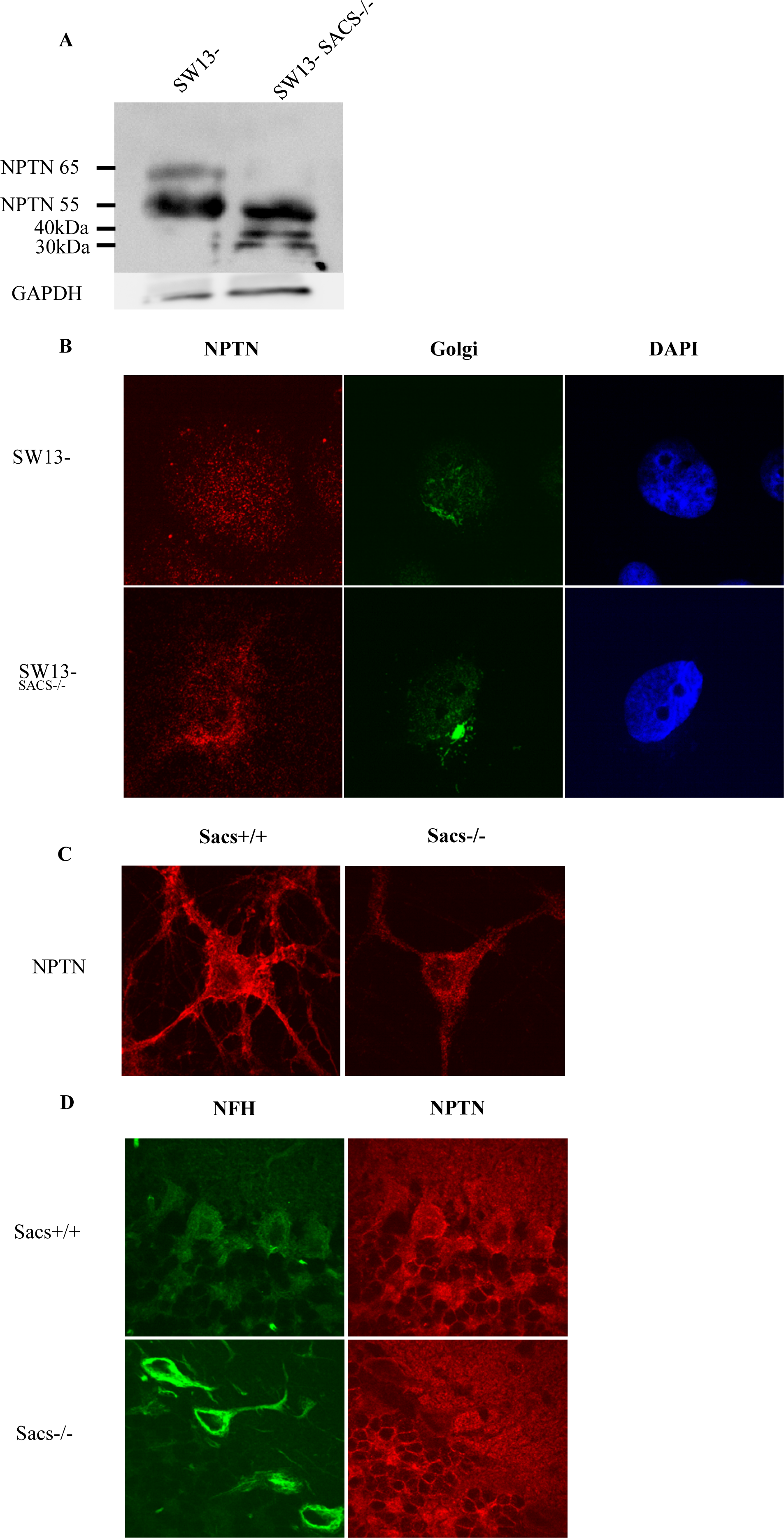
Altered electrophoretic mobility pattern of neuroplastin (NPTN) and restriction of its subcellular distribution to the soma in SW13^-*SACS*-/-^ cells. **A)** Western blot analysis showing shift in NPTN mobility in SW13^-*SACS*-/-^ cells with loss of the higher molecular weight band and appearance of lower molecular weight bands. GAPDH served as loading control. **B-C)** Representative Z-stacks of confocal images of indirect immunofluorescence labelling of NPTN and organelle markers in motor neurons of 6-week-old spinal cord cultures prepared from E13 *Sacs^+/+^* and *Sacs^-/-^* mice B) Accumulation of NPTN in the soma located in proximity to a compacted Golgi apparatus (GM130 marker) in *Sacs^-/-^* motor neurons compared to well-distributed NPTN65 and properly distributed Golgi apparatus in *Sacs^+/+^* motor neurons. **C)** Aberrant accumulation of NPTN in the soma of *Sacs^-/-^* motor neurons rather than at the membrane as seen in *Sacs^+/+^* motor neurons. **D)** Representative Z-stacks of confocal images following indirect double immunofluorescence labelling of *Sacs^+/+^* and *Sacs^-/-^* 6-months-old mice cerebellum for NPTN and NFH illustrating NF bundles and lack of membrane-localized NPTN in *Sacs^-/-^* Purkinje neurons compared to a well-distributed NF network and membrane localized NPTN65 in *Sacs^+/+^*cerebellum

## Discussion

We have identified novel interactors of the sacsin J domain that shed light on the overall functions of sacsin. The major categories of proteins pulled down from mouse brain extract were regulators of protein and organelle transport and localization, and antigen processing and presentation. While multiple Rab and associated proteins were identified, Rab1b was the most affected interacting partner in *Sacs^-/-^* tissues and cells. In the absence of sacsin, additional high molecular weight bands were present on immunoblots, suggestive of post-translational modification, and Rab1b was mislocalized, being concentrated in the soma rather than distributed throughout soma and dendrites, paralleling the compacted Golgi and ER. Rab1b abnormalities were resolved by the expression of the sacsin DNAJ or the Ubl domain, but not by other sacsin domains. DNAJ or the Ubl domain relocated Rab1b to the ER and restored the high molecular weight species of Rab1b that were lost in the absence of sacsin expression. Rab1b, like other Rab proteins, is heavily modified through geranyl-geranylation, AMPylation [31], phosphorylation, and ubiquitinylation to achieve their functions, slowing the mobility on SDS-PAGE [32]. These modifications affect localization of the protein. For example, Levin et al. demonstrated that phosphorylation modifies the membrane/cytosolic localization of Rab1b, with phosphorylated-Rab1b exclusively localizing to membranes [33]. This suggests that DNAJ and Ubl may increase Rab1b phosphorylation and promote Rab1b association with ER membrane, it is also possible that Ubl promotes the degradation of these high molecular weight proteins. Further studies will be required to identify the modifications underlying the gel-shifted species. Several post-translational modifications of Rabs have been reported including phosphorylation, ubiquitinylation, geranyl-geranylation [32, 34]. Phosphorylation of other proteins is affected in ARSACS. Hypo-phosphorylated NFH and Ataxin-2-like (ATX2L) accumulate in brain in the absence of sacsin, while stathmin (STMN1), a microtubule scaffold protein, and ADP-ribosylation Factor Like GTPase 3 (ARL3), involved in cilia formation, are hyperphosphorylated in ARSACS patient-derived fibroblasts [6, 13].

Intermediate filaments are crucial integrators of intracellular space, acting as a scaffold for organelles to regulate their positioning and dynamics [35]. This role is best demonstrated through the functional interaction between neurofilaments or vimentin IF with mitochondrial morphology [36, 37]. While IF bundling could still contribute to abnormal morphology of organelles, we demonstrated that the altered distribution of organelles and Rab1b was not secondary to the bundling of IF that is characteristic in cells lacking sacsin. The SacsJ interactors Rab1b, endophillin B2, ARF5, Rab2a, and HIP1R all have known roles in membrane dynamics. Endophilin B2 is involved in formation of endocytic vesicles and polymerization of actin around nascent vesicles [38]. Rab1b, Rab2a and ARF5 are small GTPases involved in ER-Golgi trafficking, while HIP1R, another membrane-associated protein that interacts with huntingtin, regulates clathrin-dependent endocytosis, actin polymerization, and dendritic development and synapse formation [38-40]. This impaired trafficking of membranous organelles could be at the origin of major dysregulation of cellular events including the inappropriate targeting of key plasma membrane molecules and the formation of the ATG9 membranes during autophagy.

Neuroplastin is a key cell adhesion molecule in neuronal membranes whose expression is downregulated in several neurological disorders [41]. It promotes neurogenesis and forming functional complexes with GABA receptors, ionotropic glutamate receptors, or calcium channels like PMCA1 to regulate calcium signaling and homeostasis [42, 43]. In this study, expression of Neuroplastin was reduced in *Sac*s^-/-^ cells, with loss of a higher molecular weight species on Western blots. The presence of lower molecular weight species, likely non-N-glycosylated forms is consistent with a defect of transport from the ER to the Golgi, where N-glycosylation and maturation of proteins like neuroplastin occurs. How neuroplastin contributes to ARSACS will require further studies. However, neuroplastin is highly expressed in zebrin-negative stripes of the cerebellum [44], which were described to be particularly vulnerable in *Sacs^-/-^*mice [45].

In conclusion, our investigation provides a detailed understanding of the molecular interactions and cellular processes involving the DNAJ domain of sacsin. The findings contribute to the broader knowledge of sacsin’s role in organelle dynamics, autophagy, and protein regulation, shedding light on potential implications for neurodegenerative disorders such as ARSACS.

## Declarations

the authors declare having no conflict of interest.

## Abbreviations

ARSACS: Autosomal Recessive Spastic Ataxia of the Charlevoix Saguenay
IF: intermediate filaments
NF: neurofilaments
Ubl: ubiquitin-like domain
NFL: neurofilament light polypeptide
SacsJ: Sacsin J domain
SIRPT: Sacsin Internal RePeaT domain
HEPN: Higher Eukaryote and Prokaryote Nucleotide-binding domain

