## Supplemental Material for "Interactors of sacsin’s DNAJ domain identify function in organellar transport and membrane composition relevant to ARSACS pathogenesis"

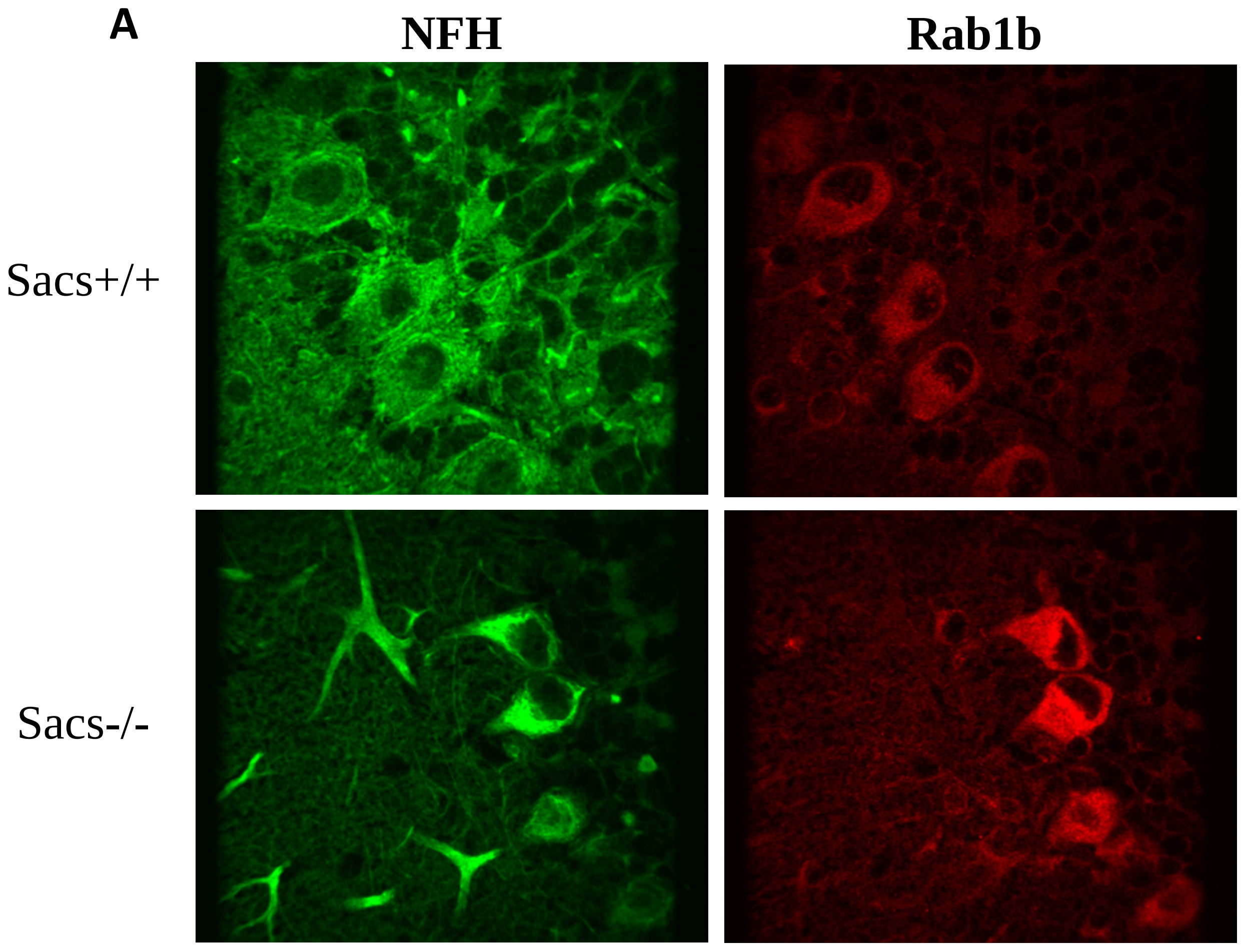


**Figure S1. Rab1b subcellular distribution is restricted to the soma and accumulates near NF bundles in *Sacs*$^{-/-}$ cerebellum neurons. A)** Representative Z-stacks of confocal images of indirect immunofluorescence labelling of Rab1b (rabbit anti-Rab1b) and NFH (mouse anti-NFH) in cerebellum of *Sacs*$^{+/+}$ and *Sacs*$^{-/-}$ mice showing accumulation of Rab1b in the soma located in proximity of NF bundles in *Sacs*$^{-/-}$ Purkinje neurons compared to well-distributed Rab1b and well-distributed NF network in *Sacs*$^{+/+}$ neurons.

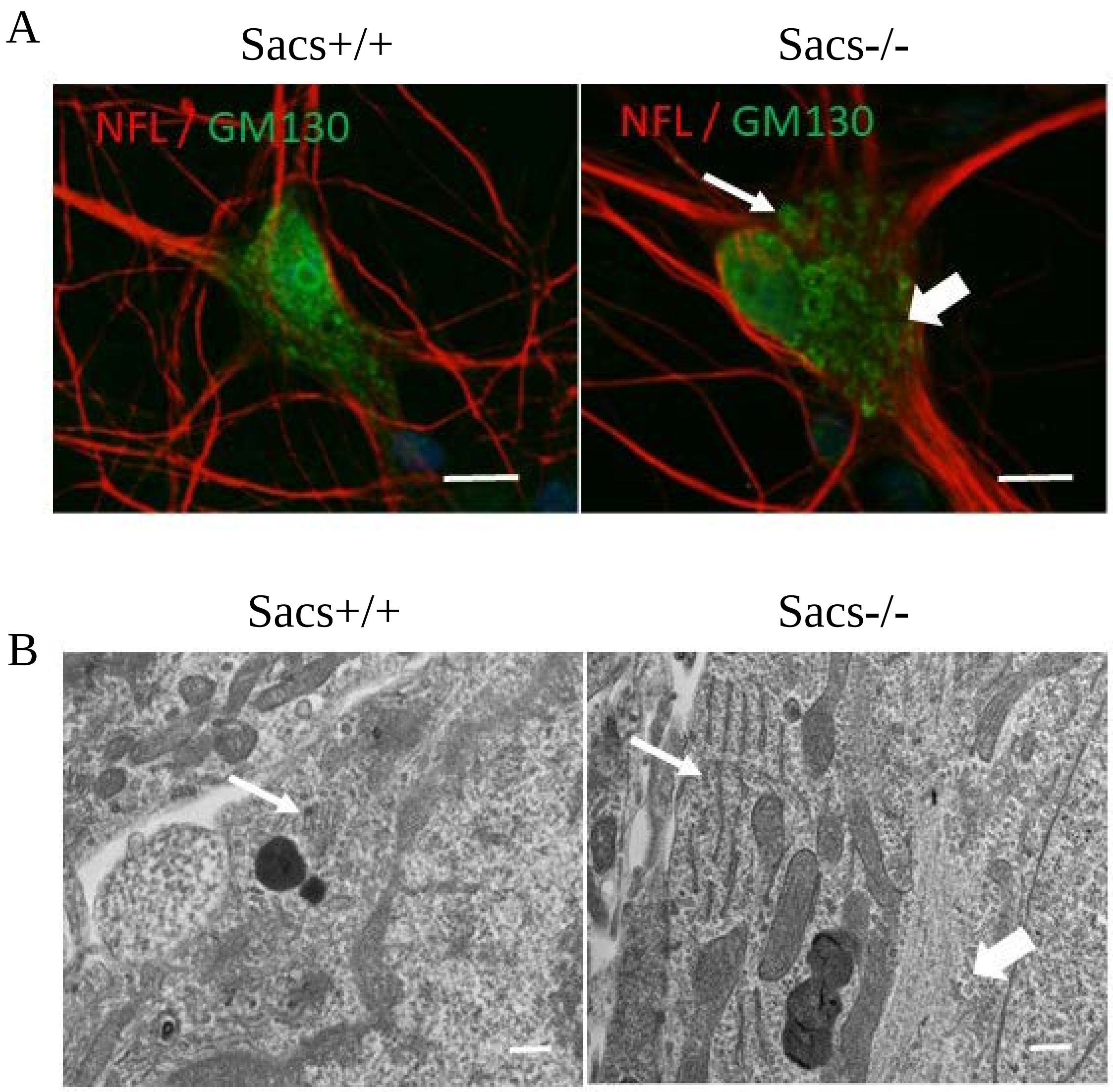


**Figure S2. Electron micrographs illustrating a neurofilament bundle in proximity to an eccentric Golgi apparatus in *Sacs*$^{-/-}$ motor neurons. A)** Representative Z-stacks of confocal images of motor neurons in 6-week-old spinal cord cultures prepared from E13 *Sacs*$^{+/+}$ and *Sacs*$^{-/-}$ mice following indirect immunofluorescence labelling for anti-GM130 (rabbit anti-GM130 in green) and anti-NFL (mouse anti-NFL in red) illustrating Golgi and neurofilament distribution. The Golgi apparatus (Right panel, thin white arrow) is displaced by neurofilament bundle (thick white arrow) in *Sacs*$^{-/-}$ motor neuron. Scale bar = 20µm. **B)** Electron micrograph *Sacs*$^{+/+}$ and *Sacs*$^{-/-}$ motor neurons showing an eccentric Golgi apparatus (thin white arrow) and a neurofilament bundle coursing in proximity (Thick white arrow) in *Sacs*$^{-/-}$ motor neurons. Scale bar (20µm).

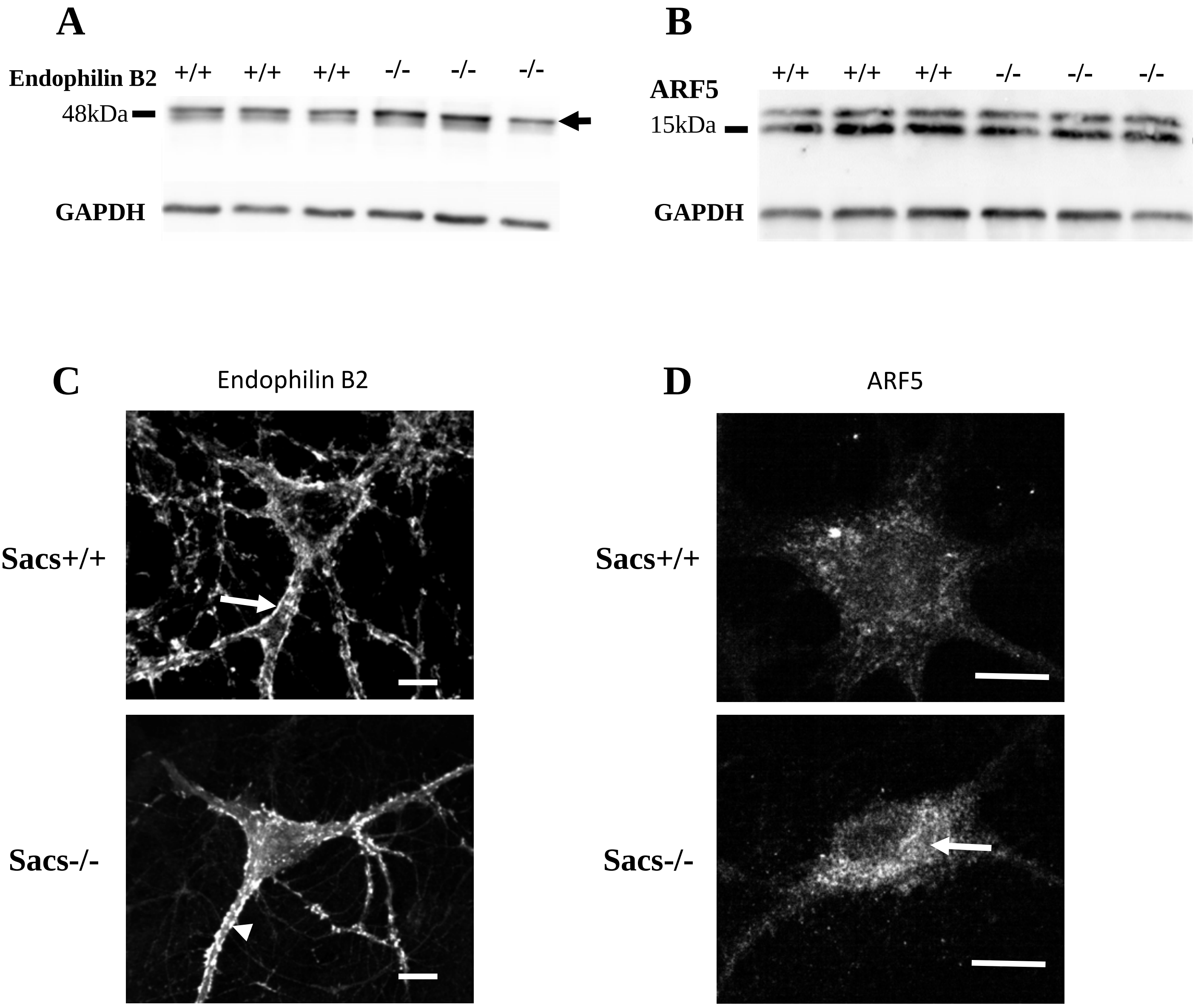


**Figure S3. Endophilin B2 and ARF5 expression and distribution are altered by the lack of sacsin. A and B)** Western blot of Endophilin B2 and ARF5, with GAPDH as loading control showing their expression levels in 6-month-old *Sacs*$^{+/+}$ and *Sacs*$^{-/-}$ mouse spinal cord. **C,D)** Representative Z-stacks of confocal images of indirect immunofluorescence labelling of motor neurons in 6-week-old spinal cord cultures prepared from E13 *Sacs+/+* and *Sacs-/-* mice for **C)** Endophilin B2 (rabbit anti-SH3GL2) and NFL (mouse anti-NFL) showing formation of abundant bulbous Endophilin B2 accumulations (arrows head) along the dendrites of *Sacs*$^{-/-}$ motor neurons compared to membrane tubulation (arrow) in *Sacs*$^{+/+}$ motor neurons (Scale bar=5µm). **D)** ARF5 (Rabbit anti-ARF5). Scale bar = 20µm.
